# Detailed analysis of chick optic fissure closure reveals Netrin-1 as an essential and conserved mediator of epithelial fusion during vertebrate embryogenesis

**DOI:** 10.1101/477729

**Authors:** H Hardy, J Prendergast, A Patel, S Dutta, V Trejo-Reveles, H Kroeger, A Yung, L Goodrich, BP Brooks, J Sowden, J Rainger

## Abstract

Epithelial fusion underlies many vital organogenic processes during embryogenesis. Disruptions to these cause a significant number of human birth defects, including ocular coloboma. We provide robust spatial-temporal staging and unique anatomical detail of optic fissure closure (OFC) in the embryonic chick, including strong evidence for roles of apoptosis and epithelial remodelling. We performed complementary transcriptomic profiling and show that *Netrin*-1 (*NTN1*) is precisely expressed in the chick fissure margin at the fusion plate but is immediately downregulated after fusion. We further provide a combination of protein localisation and phenotypic evidence in chick, humans, mice and fish that Netrin-1 has an evolutionarily conserved and essential requirement for OFC, and is likely to have a major role in palate fusion. Our data reveal that *NTN1* is a new locus for human coloboma and other multi-system developmental fusion defects, and that chick OFC is a powerful model for epithelial fusion research.

## INTRODUCTION

Tissue fusion of epithelial sheets is an essential process during normal human development and its dysregulation can result in birth defects affecting the eye, heart, palate, neural tube, and multiple other tissues^1^. These can be highly disabling and are among the most common human birth defects, with prevalence as high as 1 in 500^1–3^. Fusion in multiple embryonic contexts display both confounding differences and significant common mechanistic overlaps^1^. Most causative mutations have been identified in genes encoding transcription factors or signalling molecules that regulate the early events that guide initial patterning and outgrowth of epithelial tissues^1,3–5^. However, the true developmental basis of these disorders is more complex and a major challenge remains to fully understand the behaviours of epithelial cells directly involved in the fusion process.

Ocular coloboma (OC) is a structural eye defect that presents as missing tissue or a gap in the iris, ciliary body, choroid, retina and/or optic nerve. It arises from a failure of fusion at the optic fissure (OF; also called the *choroid fissure*) in the ventral region of the embryonic eye cup early in development^4,6,7^. OC is the most common human congenital eye malformation and is a leading cause of childhood blindness that persists throughout life^2,8^. No treatments or preventative measures for coloboma are currently available.

The process of OF closure (OFC) requires the coordinated contributions of various cell types in the fusion environment along the anterior to posterior axis of the ventral eye cup (reviewed in^4,6^). In all vertebrates studied so far, these include epithelial cells of both the neural retina and retinal pigmented epithelium, and periocular mesenchymal (POM) cells of neural crest origin^4,9–12^. As the eye cup grows, the fissure margins come into apposition along the anterior-posterior axis and POM cells are gradually excluded. Through unknown mechanisms, the basal lamina that surround each opposing margin are either breached or dissolved and epithelial cells from each side intercalate and then subsequently reorganise to form a continuum of NR and RPE, complete with a continuous basal lamina. The function, requirement and behaviour of these epithelial cells in the fusing tissue, and their fates after fusion, are not well understood.

Some limited epidemiological evidence suggests environmental factors may contribute to coloboma incidence^7,13^. However, the disease is largely of genetic origin, with as many 39 monogenic OC-linked loci so far identified in humans and the existence of further candidates is strongly supported by evidence in gene-specific animal models^4^. Most known mutations cause syndromal coloboma, where the eye defect is associated with multiple systemic defects. A common form of syndromal coloboma is CHARGE syndrome (MIM 214800) for which coloboma, choanal atresia, vestibular (inner-ear) and heart fusion defects are defining phenotypic criteria^14^. Palate fusion defects and orofacial-clefting are common additional features of CHARGE (~ 20% of cases) and in other monogenic syndromal colobomas (e.g. from deleterious mutations in *YAP1*, *MAB21L1*, and *TFAP2A*^15–17^), suggestive of common genetic mechanisms and aetiologies, and pleiotropic gene function.

Isolated (i.e. non-syndromal) OC may be associated with microphthalmia (small eye), and the majority of these cases are caused by mutations in a limited number of transcription-factor encoding genes that regulate early eye development (e.g. *PAX6*, *VSX2* and *MAF*^4,8^), implying that abnormal growth of the eye prevents correct OF margin apposition and that fusion defects are a secondary or an indirect phenotype. Indeed, none of these genes have yet been implicated with direct functional roles in epithelial fusion. However, many isolated coloboma cases also exist without microphthalmia, suggesting that in these patients, eye growth occurs normally but the fusion process itself is defective. These OCs are highly genetically heterogeneous and known loci are not recurrent among non-related patients^18^. Furthermore, despite large-scale sequencing projects, over 70% of all cases remain without a genetic cause identified^18^.

The most effective and informative models for studying OFC so far have been mouse (*Mus musculus*) and zebrafish (*Danio rerio*). Both have significant experimental advantages, including powerful genetics and robust genomic data. In particular, live-cell imaging with fluorescent zebrafish embryos has proven to be useful in revealing some intricate cell behaviours at the fissure margin during fusion^12^. However, both models are restrictive for in depth molecular investigations due to their limited temporal windows of fusion progression and the number of cells actively mediating fusion and subsequent epithelial remodelling.

Here, we present accurate staging and anatomical detail of the process of chick OFC. We show the expansive developmental window of fusion, and the sizable fusion seam available for experimentation and analysis. We take advantage of this to perform transcriptional profiling at key discrete stages during fusion and show significant enrichment for known human OFC genes, and reveal multiple genes not previously associated with OFC. Our analyses also identified specific cellular behaviours at the fusion plate and found that apoptosis was a prominent feature during chick OFC. Furthermore, we reveal Netrin-1 as a novel mediator of OFC that is essential for normal eye development in evolutionarily diverse vertebrates, and which has a specific requirement during fusion in multiple developmental contexts. This study presents the chick as a powerful model system for further OFC research, provides a novel candidate gene for ocular coloboma, and directly links epithelial fusion processes in the eye with those in broader embryonic tissues.

## RESULTS

### OFC in the chick occurred within a wide spatial and temporal window

The eye is the foremost observable feature in the chick embryo and grows exponentially through development (Fig.1a, Supplemental Fig.S1). The optic fissure margin (OFM) was identifiable as a non-pigmented region at the ventral aspect of the eye that narrowed markedly in a temporal sequence as the eye increased in size (Fig.1a). To gain a clearer overview of gross fissure closure dynamics we analysed a complete series of resected flat-mounted ventral eye tissue from accurately staged embryos at Hamburger Hamilton stages (HH.St) 25 through to HH.St34 (*n* >10 per stage; Fig.1b and Supplemental Fig.S1b). The OFM was positioned along the anterior-posterior (A-P) axis of the eye, from the pupillary (or collar) region of the iris to the optic nerve (Fig.1b). Progressive narrowing of the OFM was observed between HH.St27 to HH.St31, characterized by the appearance of fused seam in the midline that separated the non-pigmented iris from the posterior OFM (Fig.1b). Both these latter regions remained unpigmented throughout development and we found they were associated, respectively, with the development of the optic nerve and the pecten oculi - a homeostasis-mediating structure that extends out into the vitreous from the optic nerve head and is embedded in the posterior OFM (Supplemental Fig.S1)^19^. The anterior region of the pecten is attached to blood vessels that invade the eye globe through the open region of the iris OFM. This iris region remains open and the blood vessels and pecten remain throughout development and well after hatching (Supplemental Fig.S1). A recent study reported that the posterior chick OFM closes via the intercalation of incoming astrocytes and the outgoing optic nerve^20^, in a process that does not reflect the epithelial fusion processes seen during human optic fissure closure (e.g. mediated by epithelial cells of the RPE and neural retina)^9^. To assess the utility of the chick as a for human OFC and epithelial fusion, we focused solely on OFC progression in the anterior and medial eye.

**Figure 1.**
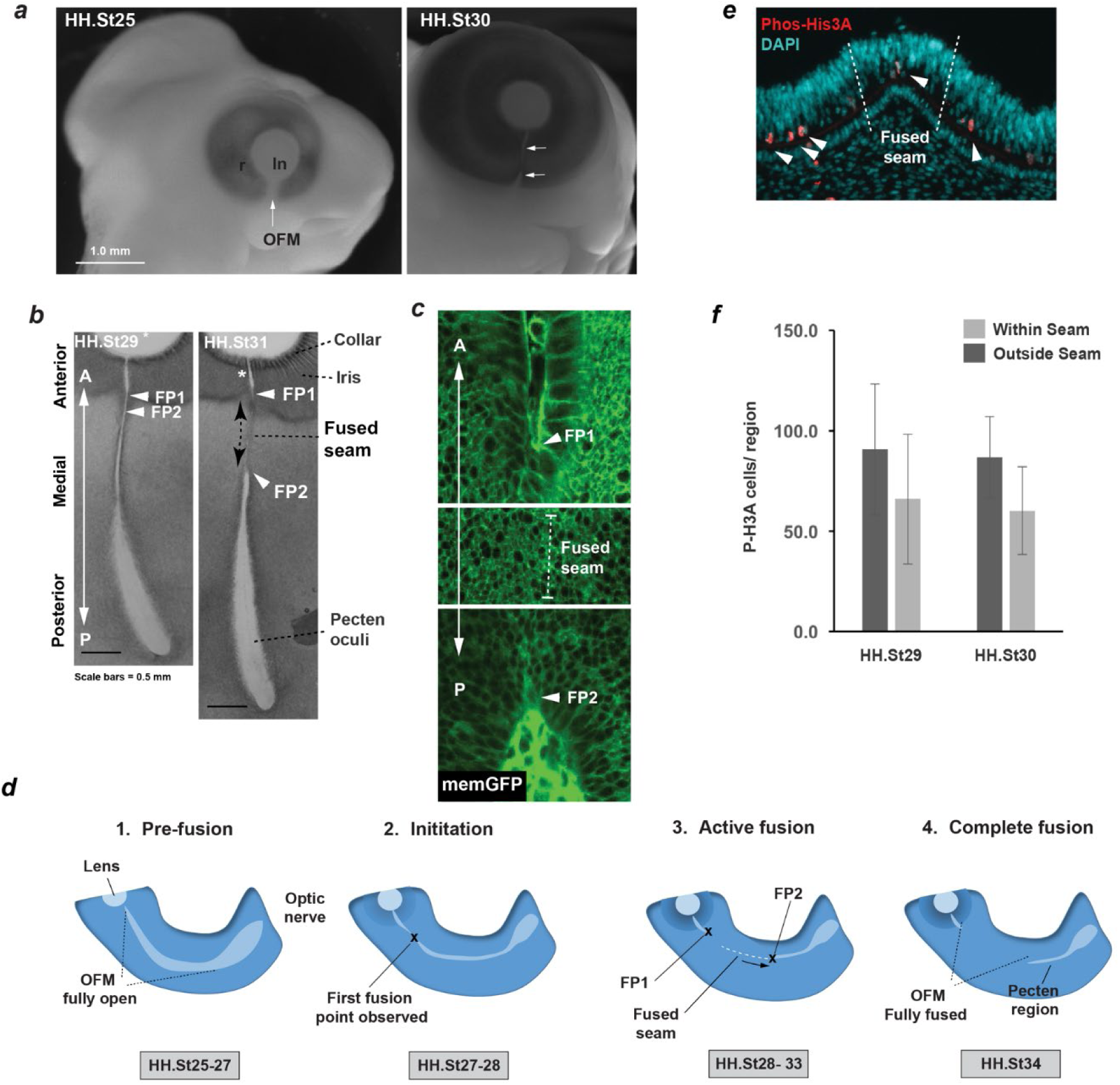
Analysis of chick optic fissure closure. (***a***) Whole mounted embryos from HH.St25, and HH.St30 illustrated the optic fissure margin (OFM; arrows) as a non-pigmented region in the ventral aspect of the developing eye. (***b***) Flat-mounted ventral eye tissues revealed fusion dynamics during closure. At HH.St29 the medial OFM had narrowed markedly along the anterior-posterior (A-P) axis between the iris and the posterior region and fusion plate 1 (FP1) and FP2 (arrowheads) are adjacently positioned in the anterior OFM. At HH.St31 the medial OFM had become fully pigmented in the fused seam, and the distance between FP1 and FP2 (arrowheads) had lengthened in the A-P axis (hatched arrows). An opening remained at the anterior region of the iris (asterisk). (***c***) Fluorescent confocal microscopy of memGFP embryos was used to unambiguously define fusion plates (arrowheads) and seam throughout all stages, supported by bright-field microscopy. All fissures were also serially-sectioned to confirm the location of each fusion plate (**Supplemental Table S1**). (***d***) Schematic representation of chick OFC progression in the anterior and medial retina. *1. Pre-fusion*: A fully open OFM is evident in the ventral retina at stages HH.St25-27; *2. Initiation*: At HH.St27-28 the first fusion is observed in the anterio-medial OFM; *3. Active fusion*: fusion extends briefly in the anterior direction but then stops in the presumptive iris to leave an open region throughout development. Fusion proceeds markedly posteriorly with FP2 extending towards the pecten. *4. Complete fusion*: Fusion stops posteriorly when FP2 meets the fused pecten region. The fusion seam is complete and epithelial remodelling occurs to generate a complete continuum of both NR and RPE. Abbreviations: ln, lens; r, retina, OF, optic fissure; A-P, anterior-posterior; FP, fusion point; HH, Hamburger Hamilton staging; RPE, retinal pigmented epithelia; NR, neural retina. (***e***) Phospho-Histone H3A immunostaining of serially-sectioned fissures at HH.St29-30 revealed the presence of mitotic cells within the apical neural retina throughout the ventral eye, but that PH3A was not enriched in the fused seam. *(**f**)* Quantified Phospho-Histone H3A immunostaining of whole-mounted fissures using confocal microscopy, confirmed that there was no significant enrichment for cell proliferation within the seam compared to equivalent nasal and temporal regions (Outside Seam) along the A-P axis.

Combining the use of confocal analyses if whole-mounted wild-type and memGFP^21^ optic fissures, and using additional histological analyses on sectioned OFMs, we unambiguously identified and measured the locations of the fusion plates and regions of fused seam along the A-P axis of the OFM (Fig.1b-c; Supplemental Fig1; **Table S1**). We observed no evidence for fusion in the medial or anterior OFM at stages before HH.St27 (Supplemental Fig.S1, **Table S1**). Fusion was first initiated between HH.St27-28 at a single point 1.3 mm (± 0.1 SD) from the pupillary collar (*n* = 10; **Supplemental Table S1**). By HH.St29, this had expanded into two fusion points FP1 and FP2 that were clearly visible (Fig.1c). FP1 had moved anteriorly to become fixed at approximately 0.5 mm (*n* = 10; range: 0.42-0.56 mm; ± 0.04 SD) from the iris collar. The position of FP1 was fixed in all subsequent developmental stages (**Supplemental Table S1**, *n* = 60 fissures analysed) and the iris region remained fully open throughout ocular development (Supplemental Fig.S1 and **Supplemental Table S1**). In contrast, FP2 displayed active movement from anterior to posterior to create a fused seam between FP1-FP2 that extended at a constant rate until HH.St34 (**Supplemental Table S1**), when FP2 was no longer distinguishable from the pecten. This expansion created a seam of 1.5 mm at its maximum length. In summary, we observed four distinct phases of fusion (Fig.1d): (1) *pre-fusion* when the entire OFM is open (up to HH.St27); (2) *fusion initiation* at HH.St27-28 in the medial OFM with the appearance of a single FP; (3) *active fusion* as two FPs separate to generate a fused seam along the A-P axis (HH.St29-33); and (4) *complete fusion* as the entire midline OFM is fused (by HH.St34). The process is therefore complete within ~ 60 hours and is active over a distance up to 1.5 mm.

### Proliferation was not a major feature of chick OFC

To determine whether the expanding seam between FP1 and FP2 was a result of active directional fusion (e.g. “zippering”), or was driven by localised cell-proliferation within the OFM seam (e.g. pushing forward static fusion plates), we used phospho-Histone-H3A (PH3A) as a marker for S-phase nuclei in mitotic cells on cryo-sections (Fig.1e) and whole-mounted (Supplemental Fig.S1) ventral eyes at HH.St29 & St30. At both stages, positive cells were located exclusively at apical regions throughout the neural retina. Analysis of whole-mounted ventral eyes also revealed the presence of PH3A cells throughout the ventral eye, but quantitative analysis using confocal microscopy of PH3A-positive cells revealed there was no significant enrichment adjacent to or in the seam (Fig.1f; Supplemental Fig.S1). This result suggests that local cell-proliferation is not a mechanism for seam expansion during chick OFC and supports a model where the fused seam expands through a zippering-like mechanism of apposed tissues.

### Chick OFC was characterised by the breakdown of basement membranes, loss of epithelial morphology and localised apoptosis

By defining the location of the fusion plates during chick OFC, we could then accurately assess the cellular environment within these regions. Immunostaining for the basement membrane (BM) basal lamina marker LamininB1 on cryo-sectioned fissure margins (Fig.2a) indicated that fusion occurs between cells of the RPE and neural retinal, as observed in human OFC^9^. Fusion between opposing margins was defined by a reduction of LamininB1 at the edges of the directly apposed fissures, followed the appearance of a continuum of BM overlying the basal aspect of the neural retina. Periocular mesenchymal cells were removed from between the fissure margins as fusion progressed. Using a histological approach, we then provided further evidence that both the RPE and NR directly contribute cells to the fusion plate (Fig.2b). We also observed that within the fusion plate there was marked epithelial remodelling of both cell types, beginning after apposition of the OFM edges. In contrast, at the fused seam (> 200μm from FP2) we observed NR and RPE cells were realigned into the apical-basal orientation and were indistinguishable from cells observed in regions outside of the OFM in the tissues, indicating that the fusion process was complete. We also noticed that pigmentation (as defined by the appearance of melanin) was not a feature of RPEin the newly-fused seam and therefore cannot be used as a superficial marker for fusion (not shown). Axonal processes were absent from fusing and fused OFM in the chick eye, as observed by Neurofilament immunofluorescence (Supplemental Fig.S2).

**Figure 2.**
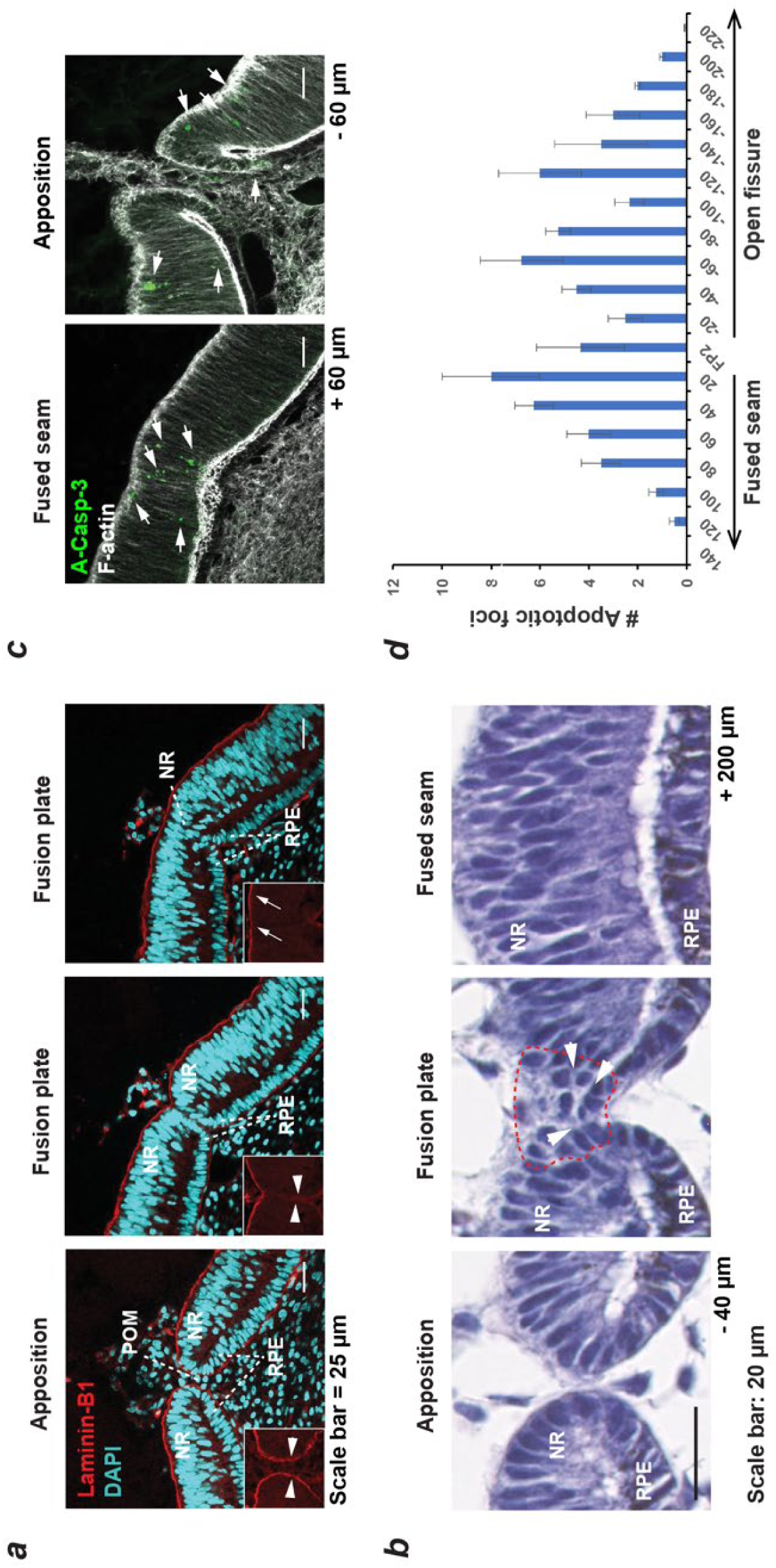
Basement membrane dynamics, epithelial remodelling and apoptosis are defining features of Chick OFC. (***a***) Immunostaining for the basement membrane (BM) component LamininB1 and nuclear staining (DAPI) illustrated that fusion was preceded by the dissolution of BM as the fissure margins came into contact at the fusion plate (arrowheads in boxes), and that fusion was characterised by the generation of a BM continuum at the basal aspect of the neural retina (arrows). Nuclear staining indicated that cells of the RPE and NR contributed to the fusion plate and that periocular mesenchymal cells were removed from the region between the apposed margins. (***b***) H&E staining Histological analysis on paraffin section at FP2 showed the fissure margins in close apposition with organised NR and RPE epithelia (−40 µm from FP2); subsequent sections showed loss of epithelial organisation *within* the fusion plate in both cell types (hatching); followed by continuous organised epithelial in both NR and RPE in the fused seam (+ 60 µm from FP2). Note that fusion occurred from contributions of both NR and RPE (arrowheads). (***c***) Immunostaining for the apoptosis marker activated Caspase-3 (A-Casp3) on cryo-sectioned OFMs (HH.St30) indicated that A-Casp3 positive foci (arrows) were localised to NR epithelia in apposed regions and in the fused seam. OFMs were counterstained with the F-actin marker Phalloidin-488 for orientation. *(**d**)* Quantification of foci from serially-sectioned OFMs at HH.St30 (*n* =4) confirmed there was significant A-Casp3 enrichment in tissues adjacent to FP2, with a graded reduction in both directions away from the fusion plate. Data in histograms are presented for the mean of all measurements; error bars = 1x standard deviations. <u>Abbreviations</u>: RPE, retinal pigmented epithelia; NR, neural retina.

Programmed-cell death has been previously associated with epithelial fusion in multiple developmental contexts but the exact requirements for this process remain controversial^1^. Even within the same tissues differences arise between species - for example, apoptotic cells are clearly observed at the mouse fusion plate during OFC^10^ but are not routinely found in zebrafish^12^. We therefore asked whether apoptosis was a major feature of chick OFC. Using HH.St29-30 eyes undergoing active fusion, we performed immunofluorescence staining for the pro-apoptotic marker activated Caspase-3. We consistently identified apoptotic foci within RPE and NR at the fusion plate, in the adjacent open fissure margin, and at the nascently fused seam with both cryo-section and whole-mount samples (Fig.2c; Supplemental Fig.S2). Foci were not found consistently in other regions of the eye or ventral retina. By quantifying the number of positive A-Casp-3 foci, we found that apoptosis was specifically enriched in the active fusion environment but was absent from fused seam >120 µm and from open regions >200 µm beyond either fusion plate (Fig.2d), indicating that apoptosis is a specific feature of OFC in the chick eye.

### Transcriptional profiling reveals genetic conservation between chick and human OFC

We took advantage of the size and accessibility of the embryonic chick eye to perform transcriptomic profiling with the objectives of: (i) assessing the utility of the chick as a genetic model for human OFCD by expression for chick orthologues of known disease genes; and (ii) to identify novel genes that are required for optic fissure closure. Using HH.st25-26 eyes (pre-fusion; approx. embryonic day E5), segmental micro-dissection of the embryonic chick eye was first performed to obtain OFM, ventral eye, dorsal eye and whole eye samples (Supplemental Fig.S3). We took care to not extract tissue from the pecten or optic nerve region of the developing OFM to ensure we obtained transcriptional data for the anterior and medial OFM only. Cognate tissues were pooled, RNA was extracted, and region-specific transcriptomes were determined using total RNAseq and analysed to compare mean transcripts per million (TPM) values (**Supplemental Table S2**). Pseudoalignment to the Ensembl chicken transcriptome identified 30,265 expressed transcripts across all tissue types. To test whether this approach was sensitive enough to reveal domain-specific expression in the developing chick eye, we compared our RNAseq expression data for a panel of genes with clear regional specific expression from a previous study of mRNA *in situ* analyses in the early developing chick eye cup^22^. Markers of the early dorsal retina *(Efnb1, Efnb2, Vsx2, Tbx5, Aldh1A1*) clustered as dorsal-specific in our RNAseq data, whereas known ventral markers (*Crx, Maf1, Pax2, Aldh6(Ald1a3), Vax1, and Rax1*) were strongly expressed in our fissure and ventral transcriptomes (Supplemental Fig.S3), which validated this approach to reveal OFC candidate genes.

We then repeated the analysis, collecting fissure, ventral tissue and whole eye and including stages HH.st26-27 (~E6; initiation) and HH.st28-30 (~E7; active fusion) as discrete time-points (Supplemental Fig.S3). Dorsal tissue was not extracted for these stages. Correlation matrices for total transcriptomes of each sample indicated one of the HH.st25-26 fissure samples as an outlier, but otherwise there was close correlation between all the other samples (Pearson’s correlation coefficient >0.9; Supplemental Fig.S3). Quantitative analyses identified 14,262 upregulated genes and 14,125 downregulated genes in the fissure margin at the three time points (Fig.3a; fissure versus whole eye. False discovery rate (FDR) adjusted *p*-value < 0.05). The largest proportion of these differential expressed genes (DEGs) were observed at HH.st25-26, most likely reflecting the periocular tissue between the fissure margins. Remarkably few DEGs were shared between stages. We used fold change (FC) analysis to identify biologically-relevant differential gene expression (Log_2_FC ≥1.5 or ≤-1) in the fissure compared to whole eye, we found 1613, 2971 and 1491 DEGs at pre-fusion, initiation, and active fusion, respectively (**Supplemental Table S3**). Refining our analysis to identify only those DEGs common across all stages revealed 12 genes with increased expression in the fissure and 26 with decreased expression (Fig.3b; Table 1). Of these upregulated fissure-specific genes, causative mutations have previously been identified in orthologues of *PAX2*, *SMOC1*, *ALDH1A3*, and *VAX1* in human patients with coloboma or structural eye malformations^4,8^, and targeted manipulations of orthologues of both *CHRDL1* and *CYP1B1* have recently been shown to cause coloboma phenotypes in *xenopus* and *zebrafish*, respectively^23,24^. The remaining fissure-specific genes (*NTN1*, *RTN4RL1*, *TFEC*, *GALNT6*, CLYBL and *RGMB*) have not been previously associated with OFC defects to the best of our knowledge.

**Figure 3.**
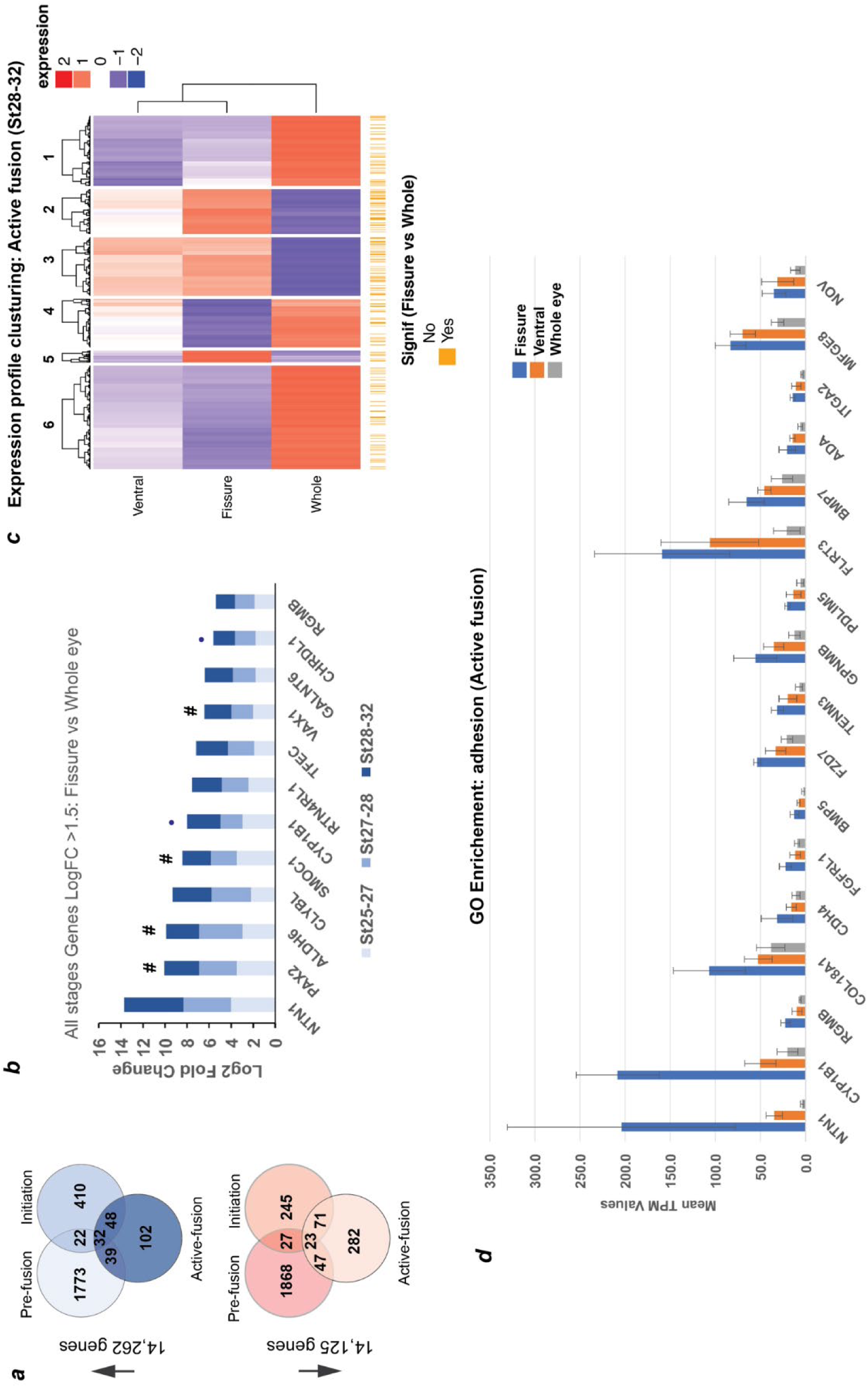
Transcriptional profiling implicates *NTN1* in chick optic fissure closure. (***a***) Transcriptional profiling using microdissected regions of the developing chick eye at E5 (HH.St25-26; Pre-fusion), E6 (HH.St27-28; Initiation), and E7 (HH.St29-31; Active fusion). revealed multiple DEGs at each stage. (***b***) *NTN1* was the highest expressing gene of 12 fissure-specific DEGs throughout all stages of chick OFC (Log2 FC >1.5; FDR < 0.05). These included the known human coloboma associated genes (indicated by #): *SMOC1*, *PAX2, VAX1* and *ALDH6*, in addition to the novel coloboma candidate (from animal studies) *CHDL1* and *CYP1B1.* (***c***) Clustering for relative expression levels at active fusion stages (HH.St28-32) revealed three independent clusters (2, 3, and 5) where expression levels matched Fissure > ventral > whole eye. (***d***) Analysis of normalised expression values (TPM) from clusters 2,3,5, at HH.St28-32 for the Gene Ontology enriched pathways (*p*< 0.0001; Biological fusion [GO:0022610], and Epithelial fusion [GO:0022610]) revealed significant fissure-specific expression for highly expressed (TPM> 100) genes *NTN1*, *FLRT*, *CYP1B1* and *COL18A1* in addition to multiple other candidates for roles in OFC. *NTN1* (TPM> 200) was the highest expressed fissure-specific gene during active fusion.

**Table 1.**
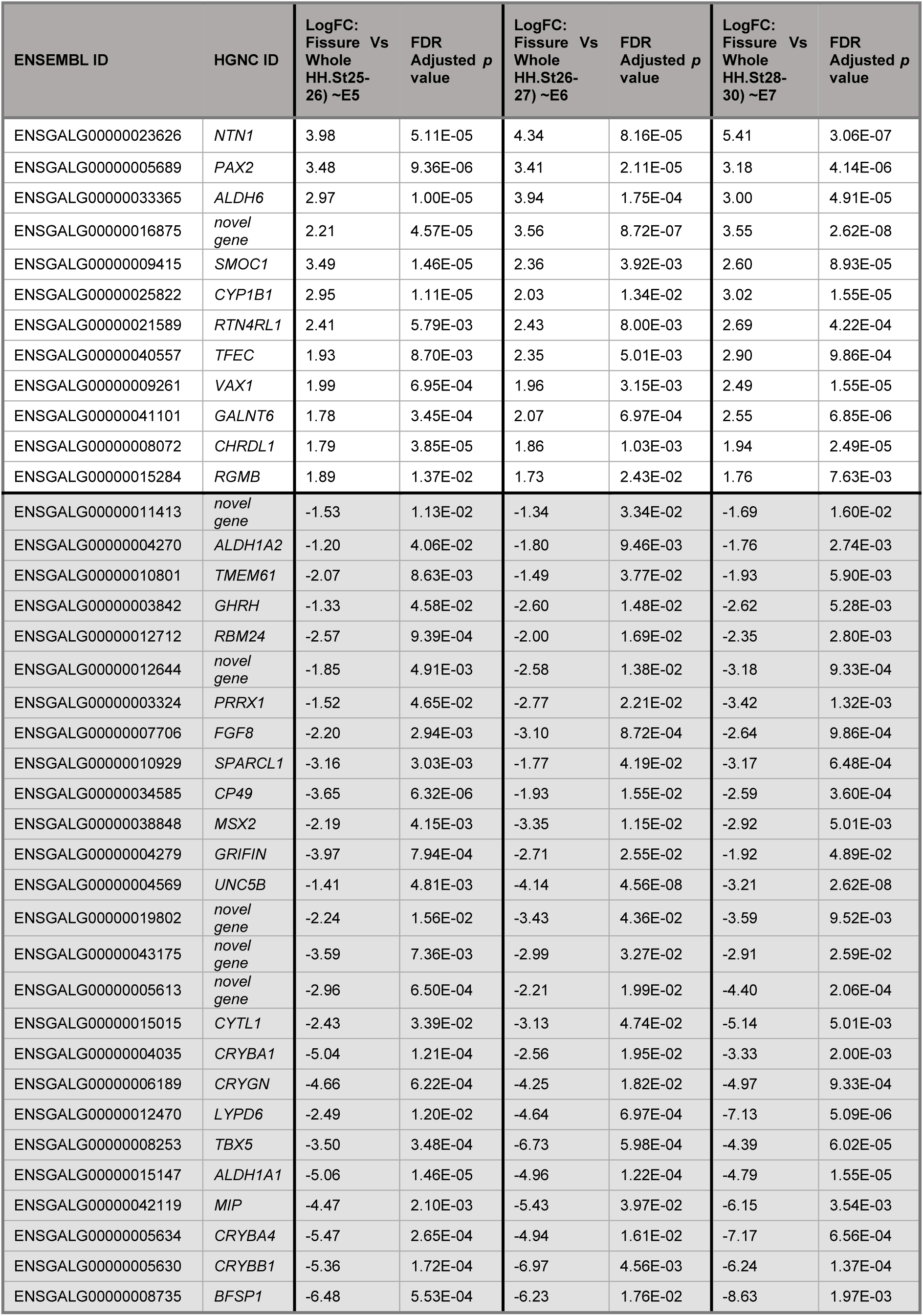
Fissure-specific DEGs (q< 0.05; LogFC: ≥1.5 and ≤-1) common to all stages analysed.

### Clustering analysis revealed *NTN1* as a fusion-specific OFC gene

Clustering for relative expression levels at active fusion stages (HH.St28-32) revealed three independent clusters (2, 3, and 5) where expression profiles matched Fissure > ventral > whole eye (Fig.3c). We hypothesised that analysis of these clusters would reveal genes with fusion-specific functions during OFC. Of the three clusters with this profile, ontology analyses showed significant enrichment for sensory organ development and eye development processes (*FDR q*< 0.001, 10 genes) and for adhesion processes (Supplemental Fig.S3; *FDR q*< 0.05, 25 genes; Biological adhesion [GO:0022610] and cell adhesion [GO:0022610]), of which 17 genes had mean TPM values >10. Within this group, multiple candidates for roles during OFC fusion were revealed, such as several transmembrane proteins, Integrin-A2, Cadherin-4, Collagen 18A1 and FLRT3 (Fig.3d). However, of these *NTN1* was the highest expressed and most fissure-specific (mean TPM values: Fissure = 204; ventral = 35; and whole eye = 4).

### *Netrin-1* is specifically and dynamically expressed in the fusing OFM

We used RNAscope and colorimetric in situ hybridisation to explore the precise location of *NTN1* transcripts in the chick eye (Fig.4a and Supplemental Fig.S4), and observed highly specific expression in both neuroepithelial and RPE cells at the fissure margins during active fusion at HH.St29-30. This was consistent to both the anterior and medial fusion plates (FP1 and FP2, respectively), and in both locations *NTN1* expression was markedly reduced in the fused seam compared to expression in the adjacent fusion plates and open margins. We then used immunofluorescence on whole-mounted OFMs at equivalent stages and observed that NTN1 protein was specifically enriched in open fissures but not in the fused seam (Fig.4b). Confocal analysis of sections from these samples revealed that NTN1 was localised to the overlying basal lamina region, in cells at the RPE-neuroepithelial junction, and at the periphery of cells at the fissure margin (Fig.4b). NTN1 was also evident in basal regions of cells within the OFM, but we were unable to determine if NTN1 was intracellular or in the ECM. Consistent with the RNAscope and transcriptomic data, NTN1 was markedly reduced at the nasal and temporal regions of the ventral retina outside from the OFM (Supplemental Fig.S4), implying that NTN1 function is fusion-specific during chick OFC.

**Figure 4.**
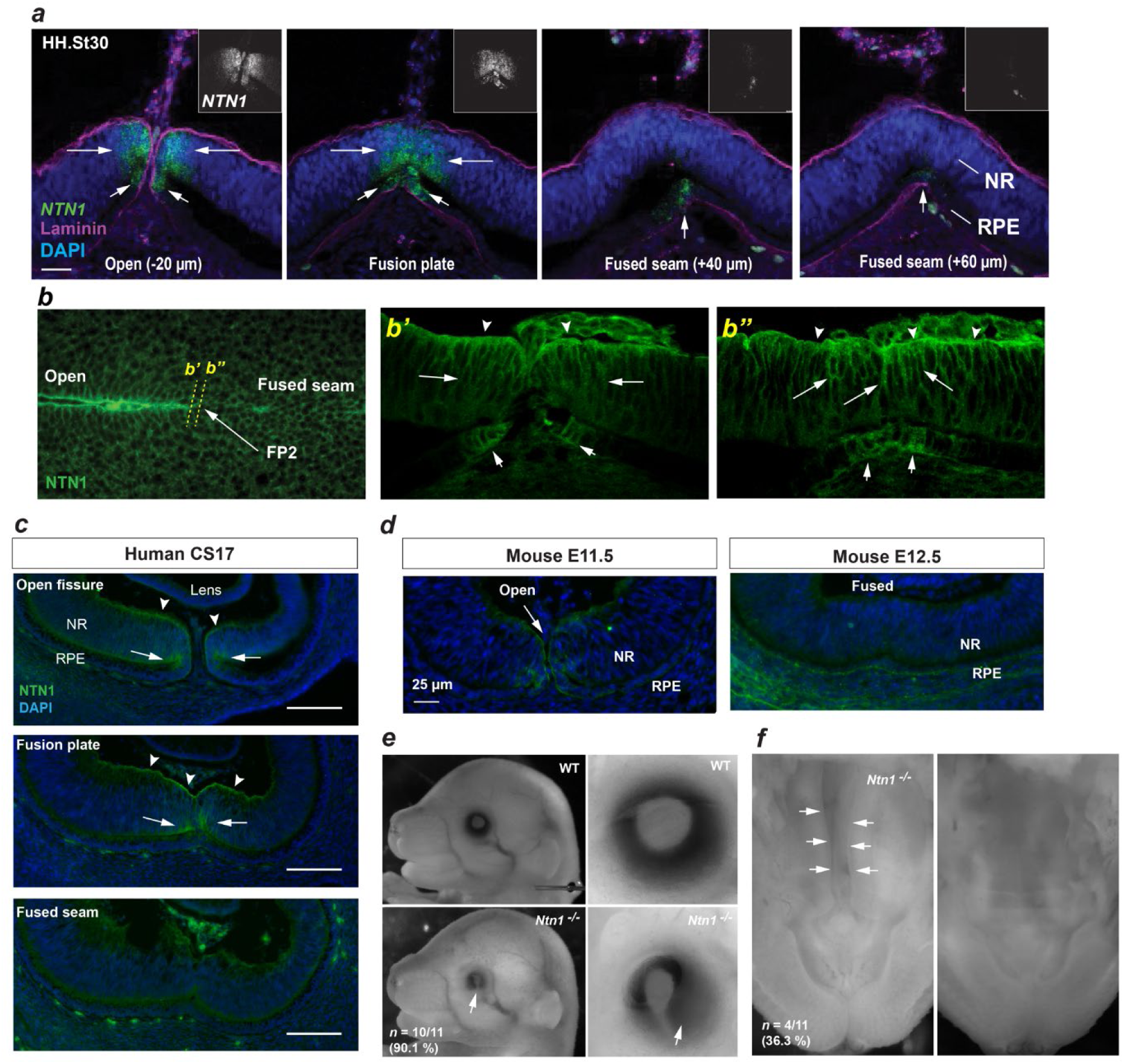
A conserved fusion-specific requirement for NTN1 in OFC and palate development. (***a***) RNAscope analysis of *NTN1* mRNA (green, and grey in insets) in HH.St29 fissures revealed dynamic *NTN1* expression (arrowheads) with strong signal observed specifically in the fusion environment at FP2 (open regions and in the fusion plate), with reduced expression seen in the adjacently fused seam. *NTN1* expression was localised to cells in both the NR and RPE (arrowheads) and fusion progression was indicated using anti-laminin co-immunofluorescence (magenta). (***b***) Confocal analysis of HH.St30 whole mount immunofluorescence for NTN1 revealed enriched protein localisation at the edges of the open fissure margins at fusion point 2 (FP2) and markedly reduced levels of protein observed in the fused OFM seam. (***b’-b”***) Cryosectioned HH.St30 fissures during fusion showed strong NTN1 immunoreactivity in the basal lamina, in regions of NR cells (dented arrowheads) at the NT (arrows) and RPE (arrowheads) at the fissure edges. (***c***) Immunostaining on CS17 human eyes revealed human Netrin-1 (hNTN1) was localised to NR epithelia (arrows) at the fissure margins and at the overlying basal lamina (dented arrowheads). Scale bar = 100 µm). (***d***) Immunostaining for mouse Netrin-1 (mNtn1) in during active fusion stages (E11.5) showed mNtn1 was localised at the open fissure margins (arrows) in the basal lamina and at the NR-RPE junction. mNtn1 was absent from there regions in fused OFM at E12.5. (***e***) Targeted *Ntn1*^−/−^ mice exhibited highly penetrant (~90 %) bilateral coloboma phenotypes (*arrows; n* = 10/11 homozygous E15.5-E16.5 animals analysed). (***f***) Cleft palate (arrows) phenotypes were observed in ~ 36 % of *Ntn1*^−/−^ embryos at E15.5-E16.5 (4/11 homozygous animals).

To test the significance of our findings to other vertebrates, we first asked whether this localisation was conserved to the human OFM. Immunofluorescence analysis for NTN1 (hNTN1) in human embryonic fissures during fusion stages (Carnegie Stage CS17) displayed remarkable overlap with our observations in chick, with protein signal localised specifically to open and fusion plate regions of OFM at the NR and RPE (Fig.4c), and an absence of hNTN1 in fused seam. Consistent with the protein localisation, RNAseq analysis on laser-captured human fissure tissue showed a 32x fold increase in *hNTN1* expression compared to dorsal eye (Patel and Sowden, unpublished data). Microarray analyses had previously observed enrichment for *Ntn1* in the mouse fissure during closure stages^25^, so we then analysed Ntn1 protein localisation in equivalent tissues in the mouse optic fissure (fusion occurs around embryonic day E11.5 and is mostly complete by E12.5^10^), and observed consistency in both cell-type and positional localisation of Ntn1 protein (Fig.4d) and that Ntn1 protein was not detected in the fused seam at E12.5 (Immunoreactivity for NTN1 was observed in the proximal optic nerve region at this stage Supplemental Fig.S4).

### Loss of Netrin causes coloboma and multisystem fusion defects in vertebrates

Our results suggested that Netrin-1 has an evolutionarily conserved role in OFC and prompted us to test if NTN1 is essential for this process. We therefore analysed mouse embryos of WT and Netrin-null (*Ntn1*^−/−^; Yung et al., 2015) littermates at embryonic stages after OFC completion (E15.5-E16.5) (Hero et al., 1998) and observed highly penetrant ocular coloboma in *Ntn1*^−/−^ mutants (> 90%; *n* = 10/11; Fig.4e). Mutant eyes analysed at earlier stages when fusion is first initiated (Hero I., 1992), were normal with fissure margins positioned directly in contact each other (*n* = 2; Supplemental Fig. S4e). We also observed variably penetrant orofacial and palate fusion defects in mutant mice (Fig. 4f; ~36%; n = 4/11 *Ntn1*^−/−^ embryos), indicating that NTN1 also has an important role in fusion during palatogenesis and craniofacial development.

We then tested whether Netrin deficiency would cause similar ocular defects in other vertebrates and generated germline loss-of-function netrin-1 mutant zebrafish (targeting both *ntn1a* and *ntn1b*) using CRISPR/Cas9 gene editing (Supplemental Fig.S4). We observed bilateral ocular coloboma in all homozygous mutant animal analysed (Supplemental Fig.S4) confirming an evolutionarily essential requirement for Netrin on OFC in diverse vertebrate species. These data revealed that *NTN1* is essential and directly required for epithelial fusion during optic fissure closure, and has an important role in palate fusion.

## DISCUSSION

### NTN1 is a novel candidate gene for coloboma and multisystem fusion defects

Our study revealed that Netrin-1 is essential for optic fissure closure in the developing vertebrate eye and is required for normal orofacial development and palate fusion. The transient and specific *NTN1* expression at the fusion plate, and the subsequent reduction/loss in fused OFM, suggests NTN1 has a direct role in the fusion process. Indeed, Netrin1-deficient mouse and zebrafish eyes both displayed highly penetrant colobomas but their fissure margins were normally apposed during fusion initiation, arguing against a broad failure of early eye development. In further support for a direct role in epithelial fusion was previously published work showing fusion failure during development of the vestibular system of both chick and mice where *NTN1*-expression was manipulated^26–28^. In this context, otic epithelia must fuse for the correct formation of semicircular canals. Taken in combination, these findings strongly implicate NTN1 as a multipotent factor specifically required for tissue fusion in multiple distinct developmental contexts. Variants near *NTN1* have been associated with cleft lip in human genome wide association studies^29,30^. While these are not monogenic disease mutations, this observation adds additional further relevance for future genetic studies of patients with coloboma. It is also consistent with our observations in *Netrin-1* knock-out animals having a high but not 100% penetrance of both coloboma and cleft lip phenotypes. Therefore, we propose that *NTN1* should be included as a candidate gene in diagnostic sequencing of patients with human ocular coloboma, and should also be carefully considered for those with other congenital malformations involving defective fusion.

### How does Netrin-1 mediate fusion

Netrin-1 is well-studied for its canonical roles in guidance of commissural and peripheral motor axons and growth-cone dynamics, with attraction or repulsion mediated depending on the co-expression of specific receptors (reviewed in^31,32^). We found that axonal processes were absent from the chick fissure margin during fusion stages (Supplemental Fig.S2), suggesting that the normal function of NTN1 may be to prevent axon ingression into the OFM to permit fusion. However, the phenotypic evidence from both the palate and vestibular system strongly support the argument that NTN1 has a non-guidance mechanistic role during optic fissure closure. Netrin orthologues have been recently associated with cell migration and epithelial plasticity in the apparent absence of co-localised canonical receptors^33–35^. In contrast, netrin acting together with its receptor neogenin combined to mediate close adhesion of cell layers in the developing terminal end buds during lung branching morphogenesis^36^. Although we observed strong *NTN1* expression in cells lining the chick OFM, and similar localisation of Netrin-1 protein in chick, human and mouse; we did not observe reciprocal expression of any canonical NTN1 receptors in our RNAseq datasets (e.g. UNC5, DCC or Neogenin; Supplemental Fig.S2). Indeed, the Netrin repulsive cue *UNC5B* was the most significantly downregulated DEG in fissure versus whole eye in our data (and from human RNAseq; Patel and Sowden, unpublished data). Therefore, it will be vitally important in future studies to elucidate the interactions of Netrin with other molecules in fusing tissues to provide insight into its mechanistic function during fusion.

### NTN1 may have a role in CHARGE syndrome

Coloboma in association with additional fusion defects of the inner ear are two of the key clinical classifications for a diagnosis of CHARGE syndrome^14^. Further phenotypes commonly associated with the syndrome are septal heart defects and orofacial clefting, both with aetiologies likely to involve fusion defects^1^. CHARGE syndrome cases are predominantly caused by heterozygous loss-of-function pathogenic variants in the chromodomain helicase DNA-binding protein 7 (*CHD7*) gene^37^. Mice lacking Chd7 display CHARGE syndrome-like phenotypes and exhibit abnormal expression of *Ntn1*^38,39^. In addition, ChIP-seq analyses have shown direct binding of Chd7 to the promoter region of *Ntn1* in mouse neural stem cells^40^. Given the amount of tissue available in the chick model, it would be possible and intriguing to confirm whether CHD7 directly regulates *NTN1* expression *in ovo* in the chick optic fissure. There is also emerging evidence that CHD7 and the vitamin A derivative retinoic acid (RA) indirectly interact at the genetic level during inner ear development^41^. Defective RA signalling also leads to significant reduction of *Ntn1* expression in the zebrafish OFM^42^, implicating a possible genetic network involving RA and *CHD7*, where *NTN1* could directly mediate developmental fusion mechanisms from these hierarchical influences.

### The chick is a powerful model for OFC

The chick is one of the oldest models for developmental biology and has provided many key insights into human developmental processes^43^. Despite this, and extensive historical study of eye development in chicken embryos the process of chick optic fissure closure has not been widely reported. Indeed, the first study appeared only recently and defined aspects of OFC at the posterior (optic nerve and pecten) region of the eye^20^, but did not describe epithelial fusion at the anterior or midline OFM. We present here the spatial and temporal sequence of chick OFC at the anatomical and molecular level, and provide strict criteria for staging the process based on a combination of broad embryonic anatomy and both ocular-and fissure-specific features (Supplemental Fig.S1). Our analyses clearly define three distinct and separate anatomical regions in the developing chick OFM: the iris, the midline region, and the pecten. Fusion initiated at the midline OFM at HH.St27/28 and continued until HH.St34, with predominantly anterior to posterior directionality. In addition, we found that OFC at the midline is a true epithelial fusion process that occurs over a large time window of approximately 60 hours, involving two fusion plates that generate up to 1.5 mm of complete fusion seam. This temporal window, the number of directly contributing cells, and the accurate staging of its progression allows unique opportunities for further experimentation. Importantly, one single fissure (from HH.St28 onwards) can simultaneously provide data for unfused, fusing, and post-fused contexts.

Our finding that chick gene expression during OFC includes multiple verified human coloboma orthologues confirms previous findings that the chick is an excellent model for human eye development and embryonic malformations^19,44,45^. These features, in combination with recent advances in chick transgenics provide a powerful toolkit to analyse cell behaviours during fusion. For example, the stable multi-fluorescent Cre-inducible lineage tracing line (the Chameleon chicken^46^) will be valuable to determine how the fissure-lining cells contribute to the fusing epithelia, while the very-recent development of introducing gene-targeted or gene-edited primordial germ cells into sterile hosts for germ-line transmission^47^ provides a rapid and cost-effective way to develop stable genetic lines to interrogate specific gene function^46,48^. Thus, our study illustrates the powerful utility of the chick as a model for investigating OFC and for the discovery of novel candidate genes for coloboma, and is perfectly timed to coincide with major new developmental biology techniques in avian systems to place the chick model as a powerful addition to OFC and fusion research.

## Summary

This study provides the first detailed report of epithelial fusion processes during chick optic fissure closure and illustrates the power of the embryonic chick eye to investigate this process further and provide insights into human eye development and broader fusion contexts.

We define the temporal framework for OFC progression, and reveal that fusion is characterised by loss of epithelial cell types and a marked increase in apoptosis. We also show that the genes expressed during chick OFC includes the specific expression of known coloboma-associated genes, and provide a broad transcriptomic dataset that may improve knowledge of candidates arising from human patient datasets. Finally, we identify that *NTN1* is specifically and dynamically expressed in the fusing fissure - consistent with having a direct role in epithelial fusion, and show NTN1 is essential for OFC and is a novel candidate for ocular coloboma and multi-fusion congenital malformations.

## MATERIALS AND METHODS

### Embryo processing

Eggs were incubated at 37°C at day 0 (E0), with embryo collection as stated throughout the text. Whole embryos were staged according to Hamburger Hamilton^49,50^. Heads were removed and either flat-mounted and imaged immediately, or placed in ice cold 4 % paraformaldehyde (PFA) in pH 7.0 phosphate buffered saline (PBS), overnight and then rinsed twice in PBS. For brightfield flat-mounts, ventral eye tissue was dissected and mounted in Hydromount (National Diagnostics HS-106) between a coverslip and glass slide, without fixation. Images were captured on a Leica MZ8 light microscope and processed using FIJI (NCBI/NIH open source software^51^).

### Immunofluorescence

For cryosections, resected ventral chick eyes were equilibrated in 15% Sucrose-PBS then placed at 37°C in 7% gelatin:15% Sucrose, embedded and flash-frozen in isopentane at −80°C. Sections were cut at 20 µm. Immunofluorescence was performed using antibodies against Laminin-B1 (DSHB #3H11; at 1:20 dilution), NF145 (Merk AB1987; 1:100), Phosphor-Histone H3A (Cell Signalling Technologies #3377; 1:200) and activated Caspase-3 (BD Pharminagen #559565; 1:400) on chick fissure sections as follows: 2x 30 min rinse in PBS, followed by 2 hours blocking in 1 % BSA (Sigma) in PBS with 0.1 % Triton-X-100 [IF Buffer 1]. Sections were incubated overnight at 4°C with primary antibodies diluted in 0.1 % BSA in PBS with 0.1 % Triton-X-100 [IF Buffer 2]. Slides were then washed in 3x 20 min PBS, followed by incubation for 1 hour with secondary antibodies (Alexa Fluor conjugated with 488nm or 594 nm fluorophores; 1:800-1000 dilution), and mounted with ProLong Antifade Gold with DAPI. Alexa Fluor Phalloidin (488 nm; Thermo-Fisher #A12379) was added at the secondary antibody incubation stages (1:50 dilution). Human foetal eyes were obtained from the Joint Medical Research Council UK (grant # G0700089)/Wellcome Trust (grant # GR082557) Human Developmental Biology Resource (http://www.hdbr.org/). For Netrin-1 immunostaining in human and mouse tissues, cryosections were antigen retrieved using 10 mM Sodium Citrate Buffer, pH 6.0 and blocked in 10 % Goat serum+ 0.2 % Triton-X100 in PBS, then incubated overnight at 4°C with primary antibody (Abcam #ab126729; 1: 300) in block. Secondary antibody staining and subsequent processing were the same as for chick. For whole-mount immunofluorescence we followed the protocol from Ahnfelt-Rønne et al^52^, with the exception that we omitted the TNB stages and incubated in IF Buffer 1 (see above) overnight and then in IF Buffer 2 for subsequent antibody incubation stages, each for 24 h at 4 °C. No signal amplification was used. Antibodies were used against Phosphor-Histone H3A (Cell Signalling Technologies #33770 at 1:1000) and Netrin-1 (R&D Systems MAB128; 1:100). Imaging was performed using a Nikon C1 inverted confocal microscope and Nikon EZ-C1 Elements (version 3.90 Gold) software. All downstream analysis was performed using FIJI. For histology and subsequent haematoxylin and eosin staining, resected eyes processed and image captured according to Trejo-Reveles et al^45^.

### In situ hybridization

RNAscope was performed on HH.St29 cryosections using a probe designed specific to chicken *NTN1* according to Nishitani et al^28^. For colourimetric in situ hybridisation, a ribprobe was for *NTN1* was designed using PCR primers to amplify a 500 bp product from cDNA prepared from chick whole embryos at HH.St28-32 (Oligonucleotide primers: Fwd 5’-ATTAACCCTCACTAAAGGCTGCAAGGAGGGCTTCTACC-3’ and Rev 5’-TAATACGACTCACTATAGGCACCAGGCTGCTCTTGTCC-3’). The PCR products were purified and transcribed into DIG-labelled RNA using T7 polymerase (Sigma-Aldrich) and used for In Situ hybridization on cryosectioned chick fissure margin tissue (as described above for immunofluorescence), and processed as described in J. Rainger’s doctoral thesis (available on request).

### Transgenic animal work

To obtain Ntn1^−/−^ mouse embryos, we performed timed matings with male and female heterozygotes and took the appearance of a vaginal plug in the morning to indicate embryonic day (E)0.5. Embryos were collected at E11.5 and E16.6 and genotyped according to Yung et al^27^. As with this previous report we observed ratios within the expected range for all three expected genotypes (28 total embryos: 13x *Ntn1*^+/−^; 10x *Ntn1*^−/−^; 5x WT – 46%; 35%; 18%, respectively). Embryos were fixed in 4 % paraformaldehyde overnight and then rinsed in PBS and imaged using a Leica MZ8 light microscope. Ntn -/-and C57Bl/6J animals were maintained on a standard 12hr light-dark cycle. Mice received food and water ad lib and were provided with fresh bedding and nesting daily. For zebrafish work, we designed gene-editing sgRNA oligos alleles according to previous reports^53–55^ to target both *ntn1a* and *ntn1b* to generate F1 founders as follows: *ntn1b* (target 1): 5’-GGACGATTCGGAGCTCGCCA-3’; *ntn1b* (target 2): 5’-GGTTGCAATGATAAGGATTT-3’; *ntn1a*: 5’-GGTCTGACGCGTCGCACGTG-3’. We then crossed *ntn1b*^−/−^;*ntn1a*^+/−^ F1s to obtain *ntn1b*^−/−^;*ntn1a*^−/−^ embryos. In these crosses, we observed coloboma phenotypes in 4/28 offspring, giving approximately 50% penetrance (7/28 expected at 100 % penetrance) (Supplemental Figure S4 and ***Supplementary Data File 1***). All experiments were conducted in agreement with the Animals (Scientific Procedures) Act 1986 and the Association for Research in Vision and Ophthalmology Statement for the Use of Animals in Ophthalmic and Vision Research.

### Transcriptional profiling

For RNA seq analysis, we carefully dissected regions of (i) fissure-margin, (ii) ventral eye, and (iii) dorsal eye, and (iv) whole eye tissue from ≥10 individual embryos for each HH stage range (Supplemental Figure S4). Samples were collected and pooled for each tissue type, and total RNA was extracted using Trizol (Thermo Scientific). Samples were collected and pooled for each tissue type and stage to obtain 3 (*n*=3) replicate RNA pools per tissue type per stage. Whole-transcriptome cDNA libraries were then prepared for each pool following initial mRNA enrichment using the Ion RNA-Seq Core Kit v2, Ion Xpress RNA-Seq Barcodes, and the Ion RNA-Seq Primer Set v2 (Thermo Scientific). cDNA quality was confirmed using an Agilent 2100 Bioanalyzer. Libraries were pooled, diluted, and templates were prepared for sequencing on the Ion Proton System using Ion PI chips (Thermo Scientific). Quantitative transcriptomics was performed using Kallisto psuedoalignment to^56^ the Ensembl (release 89) chicken transcriptome. Kallisto transcript counts were imported into R using tximport^57^ and differentially expressed transcripts identified using Limma^58^. Genes not expressed in at least three samples were excluded. To identify the relationships between samples, Log2 transformed counts per million were then calculated using edgeR^59^ and Spearman’s rank correlation was used to identify the similarities in genome-wide expression levels between samples. All RNAseq data files are submitted to the NCBI Gene Expression Ominibus database (http://www.ncbi.nlm.nih.gov/geo) with the accession number GSE84916.

## ACKNOWLEDGEMENTS

JR received funding from a BBSRC Institute Strategic Program Grant: ‘Blue Prints for Healthy Animals’ (BB/P013732/1) a Fight for Sight (UK) (Early Career Investigator Fellowship, grant number 1590/1591) and received a Travelling Fellowship (DMMTF-180520) from The Company of Biologists Ltd. JR also receives funding from a core BBSRC strategic grant to Roslin Institute. HH is supported by a Wellcome-University of Edinburgh Institutional Strategic Support Fund (ISSF). JCS and AP are supported by the Rosetree Trust, Great Ormond Street Hospital Children’s Charity and the NIHR Great Ormond Street Hospital Biomedical Research Centre. The views expressed are those of the author(s) and not necessarily those of the NHS, the NIHR or the Department of Health. We wish to thank Jenny Chen for technical assistance with zebrafish transgenics, Megan Davey at Roslin Institute for academic discussions; David FitzPatrick at The MRC IGMM for supporting the RNAseq pilot experiments; and Richard Clark at The WTCRF in Edinburgh, and Agnes Gallagher at MRC IGMM, for RNA-sequencing.

## SUPPORTING INFORMATION

**Table S1.**

Measurements of chick OFC

**Table S2.**

Quantitative measurements for Phospho-Histone H3A immunostaining during fusion seam expansion.

**Table S3.**

Kallisto analysis of RNAseq data from segmentally dissected E5 chick eyes.

**Table S4.**

Limma analysis of RNAseq data from segmentally dissected chick eyes at all stages.

**Data File 1.**

Zebrafish *ntn1* gene-edited genotyping data.

**Figure S1.**
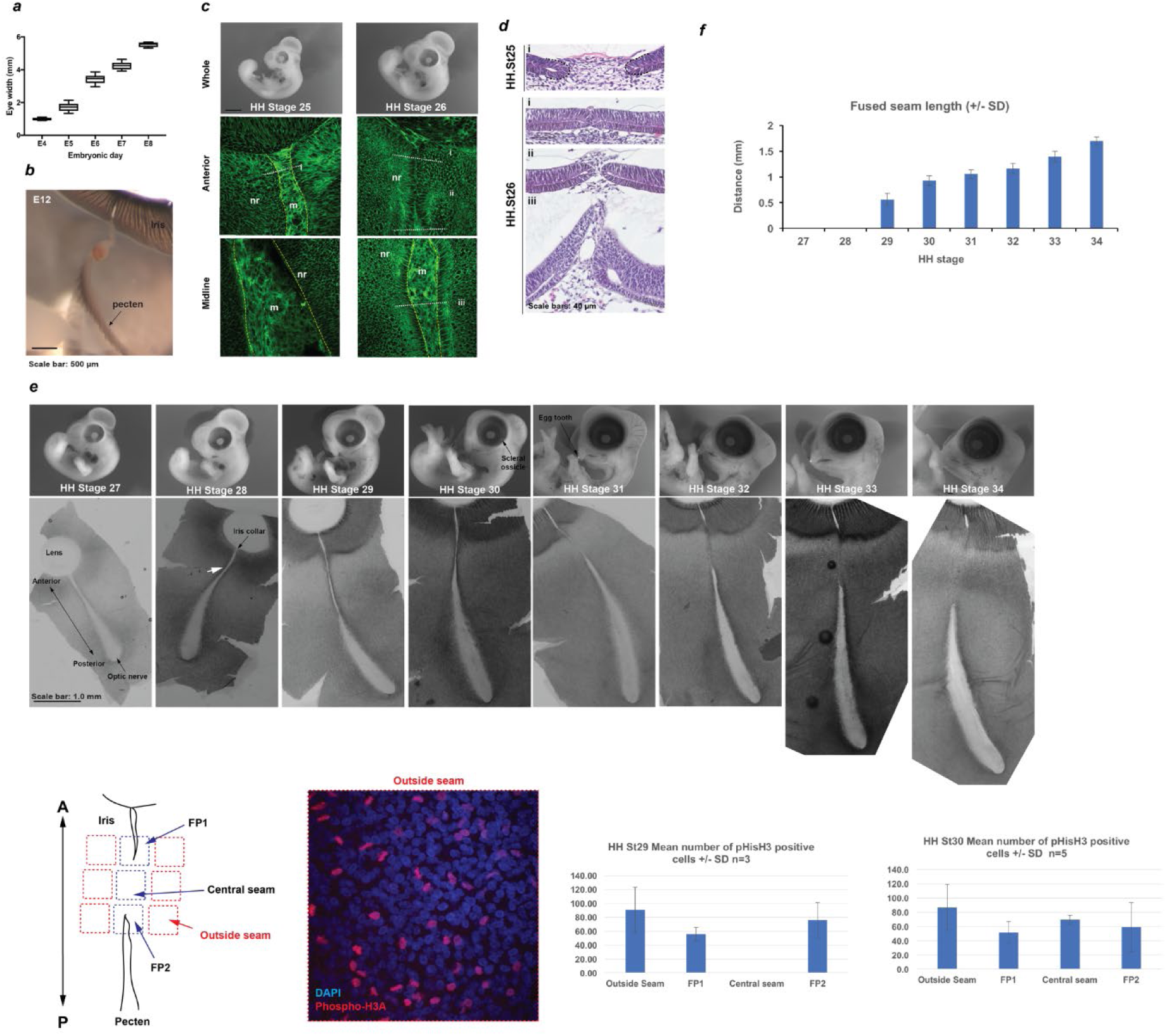
(**a**) Mean eye diameter measurements for chick embryonic days E4-E8 (*n ≥* 5 eyes per stage; one eye per embryo). (**b**) Location and orientation of the pecten oculi and associated blood vessel entering at the open iris fissure region at embryonic day 12. (**c**) Whole embryo and memGPF confocal images at HH.St25 and HH.St26 at anterior and midline illustrated the non-fused iris and medial fissure margin. (**d**) Representative H&E sections from fissures at HH.St25 and HH.St26 confirming lack of fusion at these stages. (**e**) Representative images of whole embryos and flat-mounted fissures from fusion-relevant Hamburger Hamilton^49,50^ embryonic stages. The initiating plate (white arrow) is indicated for a HH.St28 fissure. A minimum of 3 fissures were examined by confocal light-microscopy to identify fusion points and then additional samples were processed by serial cryo-sectioning to confirm fusion plates and fused seams (**Supplemental Table S1**). (***f***) Histogram to illustrate the entire fused seam length at each HH stage. (***g***) *Left*: Schema for quantifying PH3A foci within whole mounted fissures using a grid system, with a representative image showing positive nuclei. A-P axis is shown. *Right*: Histograms indicating number of PH3A-positive foci in HH.St29 and HH.St30 ventral eyes from whole mount Phospho-Histone H3A immunostaining to compare the fusion seam with non-seam regions in the ventral eye. Data shows mean PH3A+ cells per region at each stage. *Note:* seam length was too small to quantify in HH.St29 fissures, however PH3A foci were fewer at the fusion points compared to non-seam regions. Data shown is the mean of three fissures per stage with standard deviations indicated (error bars = 1x s.d.).

**Figure S2.**
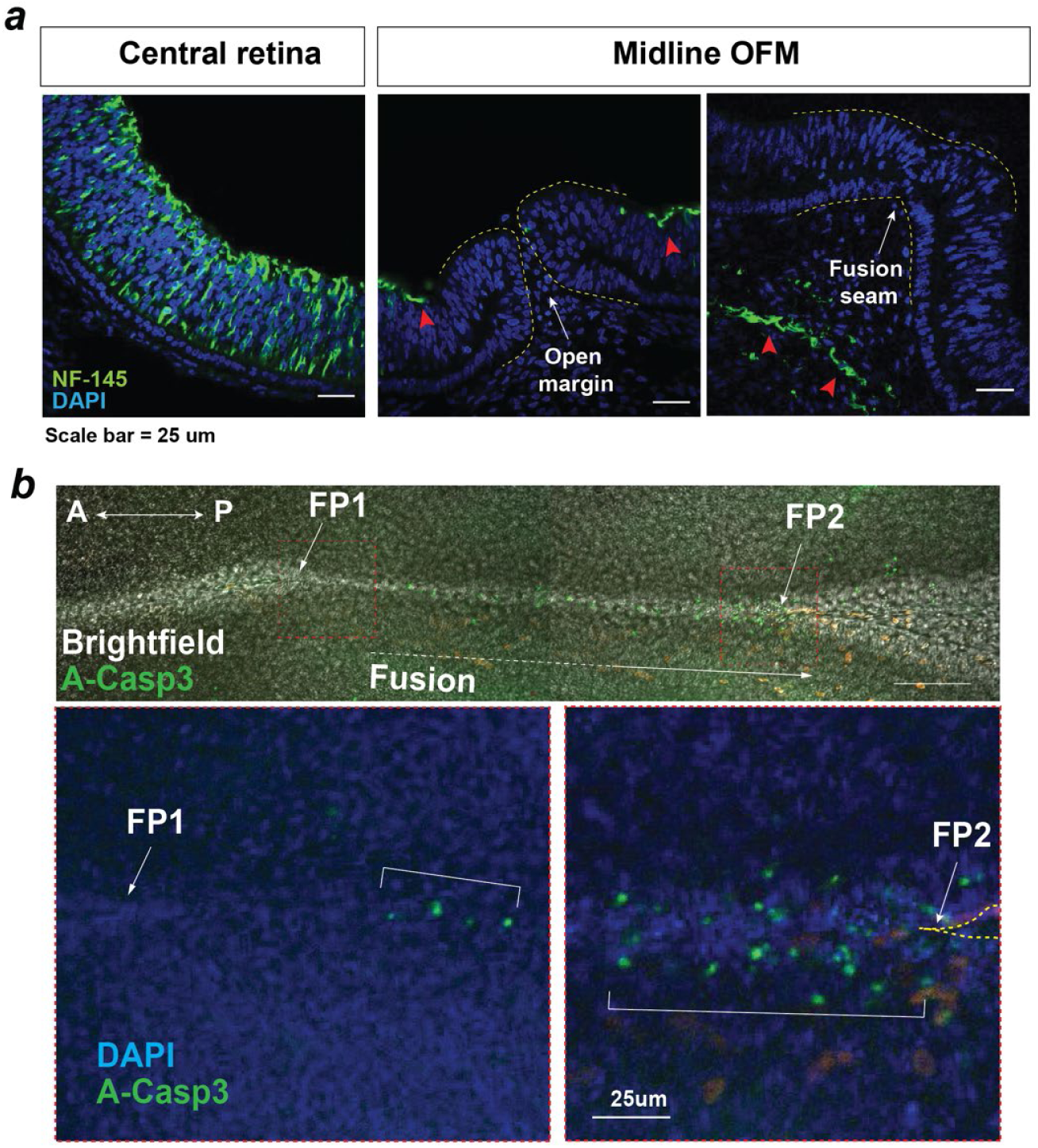
(**a**) Anti-Neurofilament medium (NF145) immunostaining showed neuron axonal processes are not a feature of OFC or present in the nascent fusion seam (HH.St30 shown). NF145 immunoreactivity (arrowheads) was observed throughout the central retina, at the ventral inner retina away from the OFM in the medial eye, and in the periocular mesenchymal regions in the fused seam. (***b***) *Top:* Confocal bright-field and fluorescence imaging of whole mount anti-activated Caspase-3 immunostaining in a HH.St30 fissure showed limited positive foci in the fused seam at a distance from the static FP1, but increased foci at the dynamic FP2. (*Below*) Enlarged images of FP1 and FP2 from (***b***) counterstained with DAPI highlighted the enrichment for A-Casp3 foci at FP2 (open margin is indicated by hatching).

**Figure S3.**
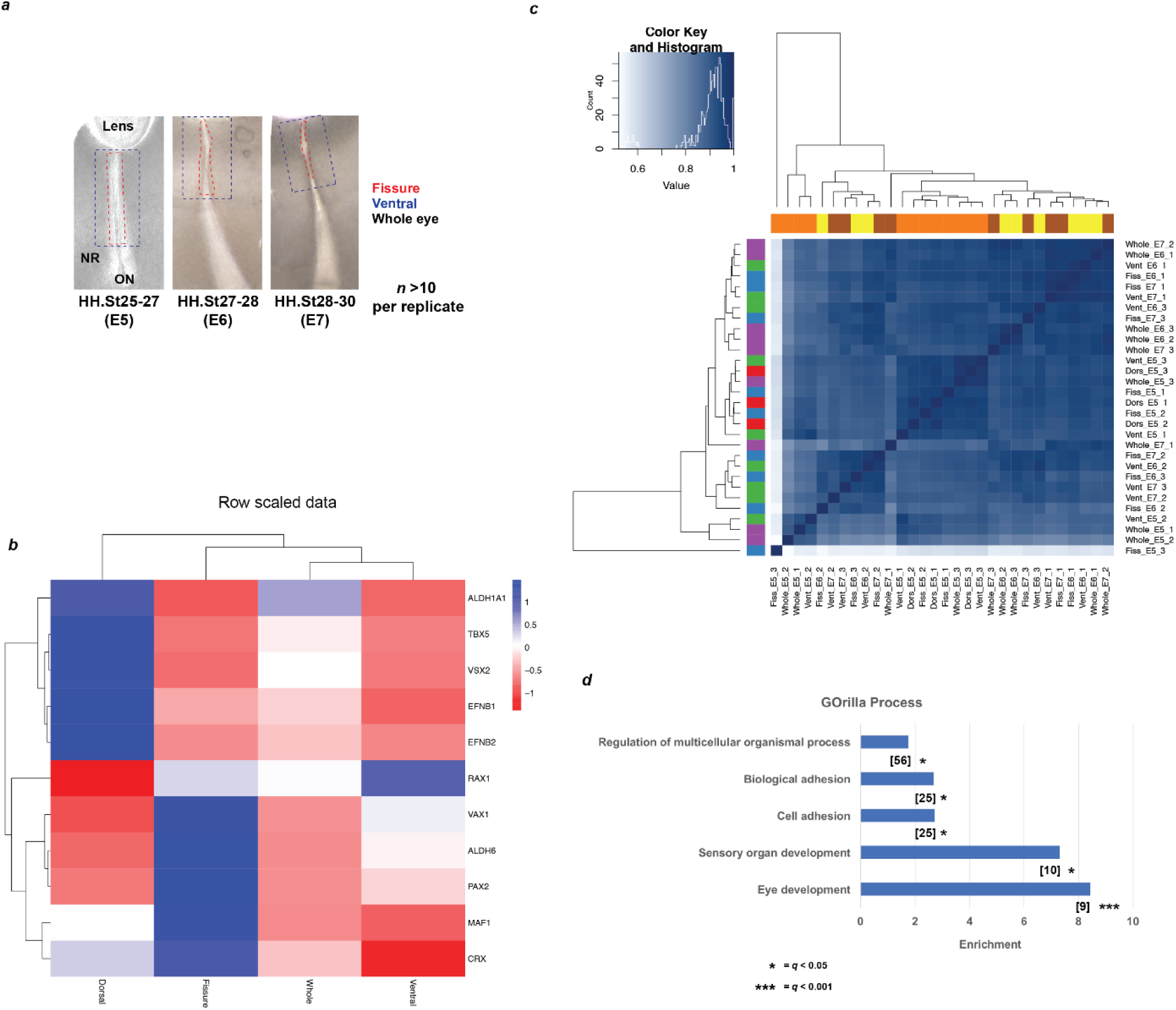
(**a**) Schema for segmental microdissection of samples prior to RNA extraction and processing for RNAseq. At least 10 fissures from independent chick embryos were used per sample, per stage. Care was taken to exclude capture of the proximal pecten region. Lenses were included for whole eye samples, for which 4 whole-eyes from independent embryos were used. (**b**) Heat map for E5 (HH.St25-27) RNAseq data shows strict domain specificity for genes with known spatial restriction^22^. (***c***) Heatmap showing a correlation coefficient of >0.9 (Spearman’s rank correlation) for genome-wide expression levels for all RNAseq samples. Note that sample Fiss_E5_3 is an outlier. (**d**) Gene ontology analysis for “Processes” using GOrilla revealed significant enrichment for five processes. Number of genes in intersection is given in brackets, FDR q values are given for each ontology. GO terms and descriptions: GO:0007423, sensory organ development; GO:0001654, eye development; GO:0007155, cell adhesion; GO:0022610, biological adhesion, GO:0051239, regulation of multicellular organismal process.

**Figure S4.**
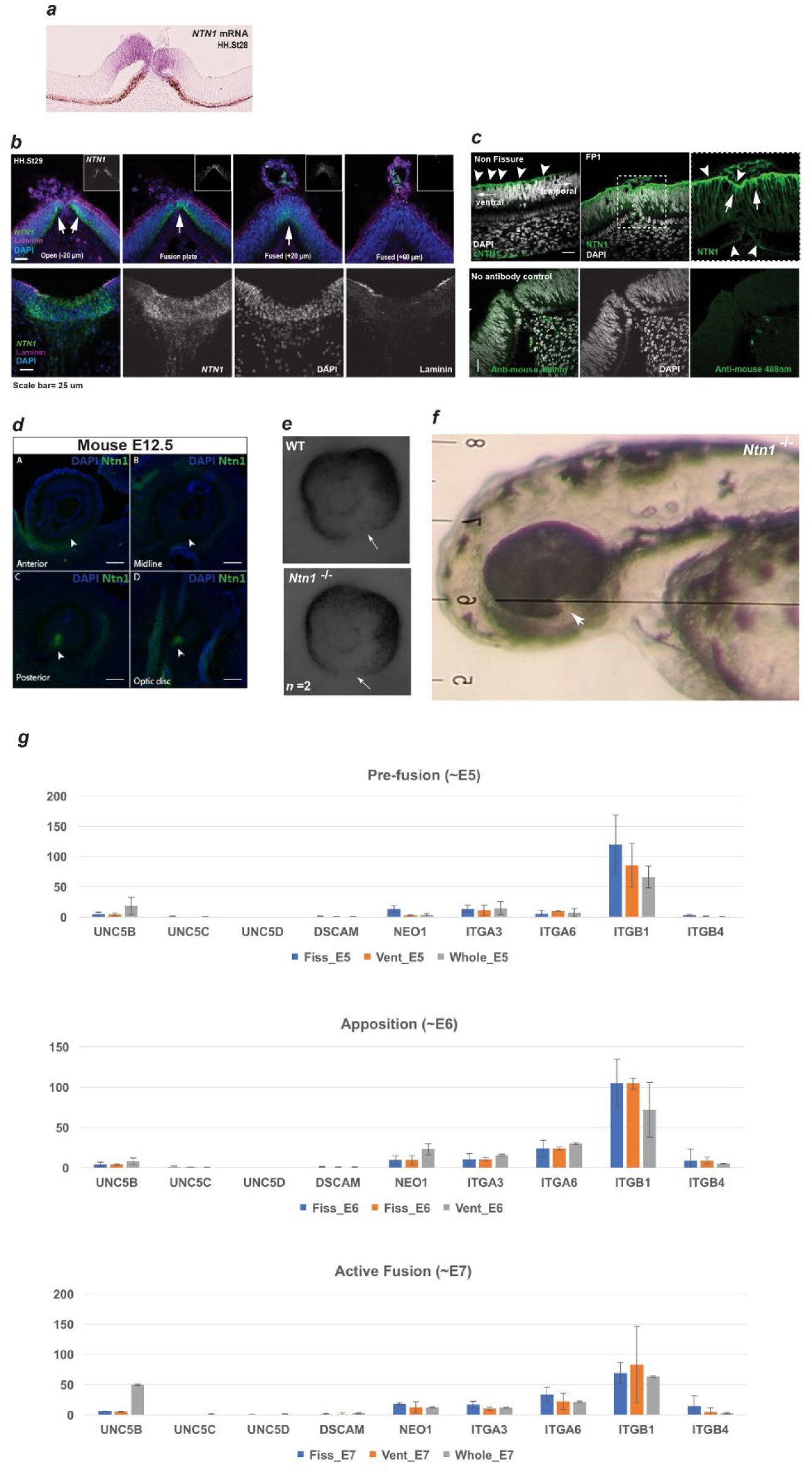
(**a**) Colorimetric *in situ* hybridisation showing the specificity of *NTN1* mRNA transcripts to the optic fissure margin in neural retina and RPE cells (HH.St28). (**b**) (*Top*) RNAscope at FP1 shows specificity to the fissure margin and graded reduction in fused seam. (*Bottom*) Positive controls for RNAscope showed strong *NTN1* mRNA signals in the basal floorplate of the neural tube at HH.St29. (**c**) Immunofluorescence analyses of NTN1 protein localisation in chick embryonic eyes illustrated positivity at the ventral retina in the basal lamina, with a gradient of reducing signal away from the fissure margin towards the temporal retina (directionality is indicated by arrows). Negative controls without primary antibody (no anti-NTN1) did not show any detectable NTN1 signal. (**d**) Immunofluorescence staining for mouse Ntn1 in E12.5 eyes post-fusion showed absence of Ntn1 signal (arrowheads) in the anterior and medially fused OFM, but persistence of Ntn1 in the posterior and optic disc regions. (**e**) E11.5 Ntn1^−/−^ embryos did not show any gross structural differences during active fusion stages and ventral tissue at the region of the fissure margins appeared to be normally apposed (Arrows). (**f**) CRISPR/Cas9 edited germline zebrafish completely lacking *ntn1a and ntn1b* exhibited fully penetrant ocular colobomas (arrow). Penetrance was calculated at 50% as coloboma was observed in 4 of 28 embryos obtained from *ntn1a*^−/+^:*ntn1b*^−/−^ crosses (7/28 expected at mendelian ratios for 100% penetrance). (**g**) Analysis of TPM values from RNAseq data at all three stages did not detect significant levels of expression for canonical NTN1 receptors in the ventral eye or fissures during OFC stages. *ITGB1* showed the highest expression values throughout all stages, but was not specific to the fissure margin or ventral eye tissues.

